# Hif1α is required for Wnt regulated gene expression during *Xenopus tropicalis* tail regeneration

**DOI:** 10.1101/2021.05.19.444777

**Authors:** Jeet H. Patel, Preston A. Schattinger, Evan E. Takayoshi, Andrea E. Wills

## Abstract

Regeneration of complex tissues is initiated by an injury-induced stress response, eventually leading to activation of developmental signaling pathways such as Wnt signaling. How early injury cues are interpreted and coupled to activation of these developmental signals and their targets is not well understood. Here, we show that Hif1α, a stress induced transcription factor, is required for tail regeneration in *Xenopus tropicalis*. We find that Hif1α is required for regeneration of differentiated axial tissues, including axons and muscle. Using RNA-sequencing, we find that Hif1α and Wnt converge on a broad set of genes required for posterior specification and differentiation, including the posterior *hox* genes. We further show that Hif1α is required for transcription via a Wnt-responsive element, a function that is conserved in both regeneration and early neural patterning. Our findings indicate a regulatory role for Hif1α in Wnt mediated gene expression across multiple tissue contexts.

## Introduction

Following severe injury, all organisms initiate a wound healing program, though regenerative outcomes vary across species (Kakebeen & Wills, 2019). Organisms such as planaria, axolotls, and zebrafish exhibit complete and robust tissue regeneration, being able to replace lost limbs and organs with a variety of cell types and proper structural organization (Erickson & Echeverri, 2018; Ivankovic et al., 2019; Kakebeen & Wills, 2019; Marques et al., 2019). Conserved injury-induced stresses including immune cell activation, inflammation, and reactive oxygen signaling occur in both regenerative and non-regenerative species but are essential for proper regeneration in species with this capability (Kakebeen & Wills, 2019; Phipps et al., 2020). The downstream events that selectively link universal stress signals to regenerative growth in only some animals remain a persistent enigma in regeneration biology.

A specific gap in our understanding of stress signaling is how injury cues are coupled to changes in gene expression that direct patterning in regenerating tissues. Macrophages, reactive oxygen species (ROS), and oxygen flux at the wound edge are all injury induced stresses that are essential for regeneration in multiple species (Ferreira et al., 2018; Godwin et al., 2013; Love et al., 2013; Pirotte et al., 2015; Romero et al., 2018; Simkin et al., 2017), suggesting that transcription factors activated by these signals represent a possible link between injury and downstream gene activation. One such stress-activated transcriptional regulator is Hypoxia Inducible Factor 1α (Hif1α), which acts downstream of ROS, inflammatory signaling, and oxygen sensing, and is poised to couple cell extrinsic stressors with regenerative gene expression (Movafagh et al., 2015). Previous work has shown that Hif1α is necessary and sufficient for regeneration of *Xenopus laevis* tails (Ferreira et al., 2018). However, the downstream effects of Hif1α, in particular its transcriptional targets, are not known in a regenerative context. Moreover, while Hif1α canonically regulates apoptosis, cell proliferation, and cellular metabolism, it has also been shown to have roles in patterning and differentiation which have not been explored in this context. Hif1α is required for proper neural tube development in mice (E. Y. Chen et al., 1999; Iyer et al., 1998) and neural crest chemotaxis in both *Xenopus* and chicks (Barriga et al., 2013). Hif1α also modulates Wnt signaling to direct gene expression and establishment of neural and skeletal muscle fates (Kaidi et al., 2007; Majmundar et al., 2015; Rohwer et al., 2019; Večeřa et al., 2017).

Wnt signaling is a deeply conserved factor in regeneration and is regarded as a key component of a complete regenerative response. It has been shown to be sufficient to drive regeneration of *Xenopus* tails, posterior structures in planaria, and limb buds in chick embryos (Gurley et al., 2008; Kawakami et al., 2006; Lin & Slack, 2008). In planaria, knockdown of *wntP-1* results in ectopic anterior regeneration, suggesting that Wnt is critical in regulating posterior identity in new tissues (Petersen & Reddien, 2009). Other regeneration models, including zebrafish fin and heart, as well as *Xenopus* and axolotl limbs, have also been shown to require Wnt signaling to promote growth and cell type specification (Kawakami et al., 2006; Strand et al., 2016; Wehner et al., 2014). Work in *Xenopus* suggests that Wnt acts upstream of other developmental signaling pathways such as BMP, Notch, and FGF in establishing muscle and neural cell fates in regeneration (Beck et al., 2003; Slack et al., 2004). While Wnt signaling appears to be a key regulator of regenerative outcomes, it is not established how injury stresses are coupled to Wnt activation nor how these cues are regionally interpreted.

Here we set out to define the downstream functions of Hif1α in *Xenopus tropicalis* tail regeneration, a versatile model for complex appendage regeneration (Kakebeen & Wills, 2019; Li et al., 2016). We show that pharmacological inhibition of Hif1α with two orthogonal reagents inhibits tail regeneration in *X. tropicalis,* as in *X. laevis,* and that Hif1α is required for muscle and axon growth. We find that genes sensitive to Hif1α perturbation include posteriorizing factors known to be targeted by canonical Wnt signaling. To broadly address whether these genes are regulated by both Hif1α and Wnt, we perform RNA-sequencing on Hif1α and Wnt antagonized tails during regeneration to show that Hif1α and Wnt are required for activation of largely overlapping groups of genes. Notably, both Hif1α and Wnt are required for posterior *hox* gene expression in the regeneration bud, suggesting a unique role for Hif1α in spatial patterning in this context. We find that Hif1α is required for activation of gene expression via Wnt-responsive promoter elements (WRE) during regeneration, as well as during early neural development. Our findings suggest a model in which Hif1α facilitates expression of specific Wnt target genes by regulating activation of Wnt responsive promoters and clarify a mechanism for the transduction of injury-induced signals upstream of developmental patterning genes.

## Results

### Hif1α is necessary for muscle and axon regeneration in *X. tropicalis*

Previously, Hif1α inhibition via Echinomycin (Ech), has been shown to reduce tail regeneration in *X. laevis* tadpoles (Ferreira et al., 2018; Kong et al., 2005). We first set out to determine if Hif1α antagonism inhibited tail regeneration in *X. tropicalis* tadpoles using Ech, which inhibits Hif1α binding to DNA, as well as 2-methoxyestradiol (2ME), which inhibits Hif1α nuclear localization and transcriptional activity (Mabjeesh et al., 2003). We identified effective doses of both inhibitors which reduced tail regeneration but did not cause lethality or reduce health (Supp. Fig. 1). Compared to DMSO treated tails, tails treated with 5μM 2ME or 0.5μM Ech had significantly reduced tail lengths compared to control clutchmates 72 hours post amputation (hpa) (Fig. 1A,B) We scored regeneration by binning tails into 4 categories ranging from no regeneration to complete tail regeneration and found that Hif1α inhibited tails had significantly reduced regenerative outcomes relative to DMSO controls (Fig. 1A,C) (Beck, 2012). These data confirm that Hif1α is required for tail regeneration in *X. tropicalis.*

**Figure 1:**
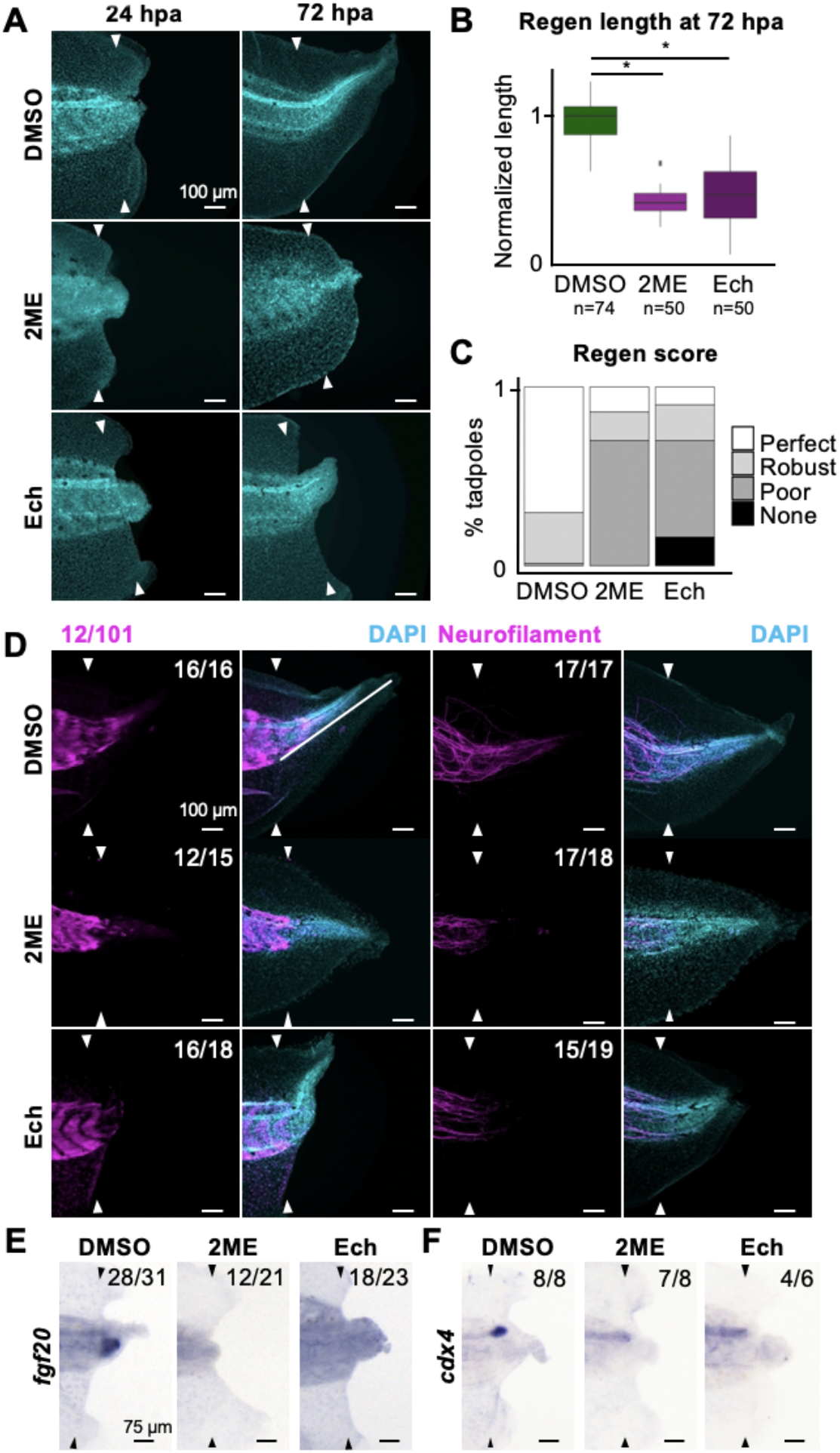
Hif1α is required for regeneration of muscle and axons. A) Dapi counterstained tails at 24 and 72hpa following treatment with DMSO, 2ME, or Ech. B) Quantification of regeneration length normalized to DMSO clutchmates. Statistical significance between groups was determined by ANOVA (p<2e-16) followed by Tukey’s posttest. * indicates p < 0.001. C) Regeneration scores binned to great (full tail regeneration), good (incomplete fin regeneration), poor (very little regeneration), or none. The treatments have statistically significant distribution of phenotypes (chisquare test, p < 2.2e-16). D) Immunohistochemical stains for 12/101 (muscle) and neurofilament (axons) at 72hpa after treatment with DMSO, 2ME, or Ech. E-F) *in situ* hybridization for *fgf20* (E) or *cdx4* (F) at 24hpa following DMSO, 2ME, or Ech treatment. Indicated numbers in D-F represent number of tails with prevented phenotype over total number assayed. Scale bars in A and D-F are 100 μm. Arrowheads in A and D-F indicate amputation plane.

We next asked if axial tissues were specifically sensitive to loss of Hif1α function. We noted that neither 2ME nor Ech completely abrogated regeneration in all treated tadpoles, but that axial tissues always appeared to be reduced (Fig. 1C). We performed immunohistochemical staining for axons and muscle at 72hpa in tadpoles treated with DMSO, 2ME, or Ech and found that Hif1α antagonism resulted in tails with little to no muscle (12/101) or axon (neurofilament) regeneration, despite a large degree of fin regeneration *(Fig. 1D).* We then asked whether Hif1α was required for the activation of neural or mesodermal fate specification genes during regeneration. To this end, we performed *in situ* hybridization for *fgf20* and *cdx4,* posterior mesoderm and neuronal markers, respectively (Lea et al., 2009; Northrop & Kimelman, 1994). We found that at 24hpa, both *fgf20* and *cdx4* are strongly induced in the regeneration bud in controls but not if Hif1α is inhibited (Fig. 1E,F). These results suggest that Hif1α might be specifically required for the activation of posterior fate genes during regeneration.

### Hif1α and Wnt are necessary for expression of similar genes during regeneration

Having found that Hif1α is required to activate *fgf20* and *cdx4,* known Wnt target genes, we hypothesized that Hif1α might broadly target the same genes as Wnt signaling (Chamorro et al., 2005; Haremaki et al., 2003). To test the requirement of Wnt signaling in regeneration, we used the Wnt inhibitor IWR-1 (IWR), which stabilizes the axin destruction complex (Borday et al., 2018; Chen et al., 2009). We find that treatment with 10μM IWR greatly reduces regeneration length and quality, in agreement with previous studies inhibiting Wnt in *Xenopus* tail regeneration (Supp. Fig. 1) (Lin & Slack, 2008). We then performed RNA-sequencing to identify regeneration-induced changes in gene expression dependent on either Hif1α or Wnt. We collected tail tissue 250μm anterior to the amputation plane at either 0 or 24hpa following treatment with DMSO, 2ME, Ech, or IWR (Fig. 2A). RNA-Seq libraries were prepared and analyzed in triplicate. When multidimensional scaling (MDS) was applied to visualize the relationship between groups (Robinson et al., 2010) we found that biological replicate libraries clustered tightly together, demonstrating their reproducibility (Fig. 2B, Supp. Fig. 2A). We also find that 0hpa tails are largely distinct from all four 24hpa groups (Supp. Fig. 2A). To visualize gene expression across treatments, we normalized to DMSO and generated a heatmap. This revealed that the directionality of changes in expression relative to DMSO were largely conserved following inhibition of Hif1α or Wnt (Fig. 2C, Supp. Fig. 2B), with 1443 genes downregulated by all treatments (Fig. 2D). These results suggest that Hif1α and Wnt direct similar gene expression programs during regeneration and identify genes regulated by both signals. PANTHER Gene Ontology (GO) Enrichment analysis on these shared, down-regulated genes called terms associated with several known Hif1α regulated processes, including erythrocyte cell homeostasis, as well as several not commonly associated with Hif1α, including macromolecule biosynthesis and mRNA splicing (Table S1). Notably, developmental pattern specification terms affiliated with Wnt signaling were called (Fig. 2E). This result, together with our observed loss of posterior markers *fgf20* and *cdx4* (Fig. 1E,F), suggested that Hif1α may be a transcriptional activator of posterior patterning genes in the regenerating tail.

**Figure 2:**
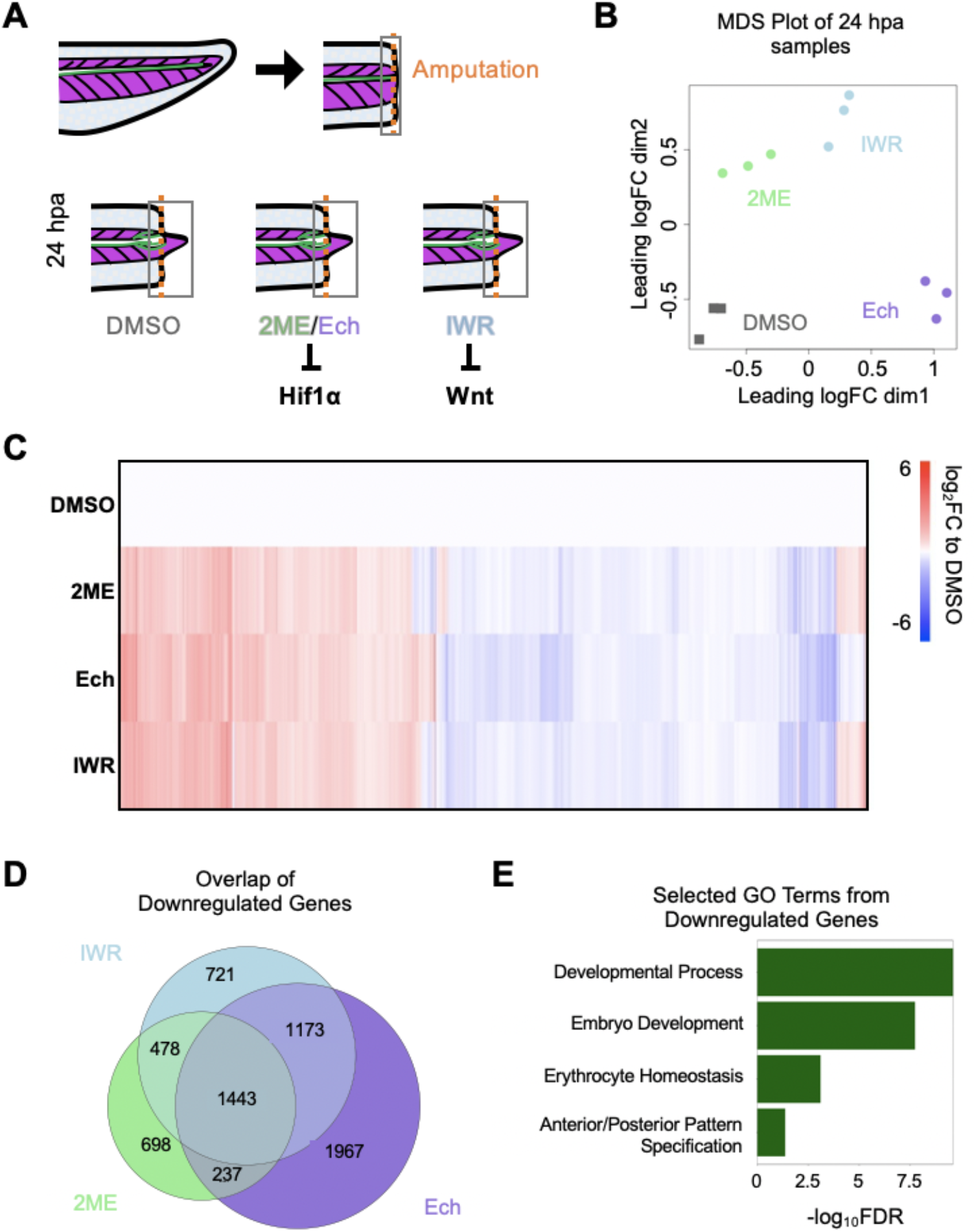
RNA-sequencing reveals shared gene regulatory roles for Hif1α and Wnt. A) Schematic depicting experimental setup for RNA-sequencing. Tails were amputated and treated with DMSO, 2ME, Ech, or IWR and collected at 0 or 24hpa for sequencing. Purple chevrons in tails represent somites and green line indicates spinal cord. Amputation plane is marked with an orange dotted line. B) MDS plot depicting relationship between samples. C) Heatmap of gene expression as a log_2_FC relative to DMSO clustered by expression. *Lower bound of color scale represented values from −6 to −15. D) Venn diagram showing overlap in downregulated genes (FDR < 0.05 and log_2_FC < −0.2). E) Selected GO terms from PANTHER gene ontology enrichment plotted by -log_10_FDR.

### Posterior *hox* gene expression in the regeneration bud requires Hif1α and Wnt

Noting that Hif1α and Wnt inhibition resulted in a downregulation of genes involved in patterning, we examined the genes under these GO terms and noticed that more than half of all *hox* genes were downregulated under these treatments (Fig. 3A,B). We examined *in situ* expression patterns at stages 32-34 on Xenbase of *hox* genes which were differentially expressed in all 3 of our treatment groups to determine the normal expression of this family of patterning related genes (Bowes et al., 2010). Notably, all but 1 of the *hox* genes that were downregulated following Hif1α and Wnt inhibition have expression in the most posterior region of tailbud stage tadpoles. The only gene that was upregulated in each group was *hoxb3,* which we noted had a distinct domain in the anterior of the tadpole (Fig. 3C). Using previously published RNA-sequencing data over a regeneration timecourse in *X. tropicalis* (Chang et al., 2017), we asked what the normal expression dynamics of the *hox* genes were across a regenerative timecourse. Relative to uninjured tadpoles, the expression of most *hox* genes is decreased at 0 and 6hpa but then increased by 15 and 24hpa (Fig. 3D). Of those genes that are activated during regeneration, most fail to increase in expression when Hif1α and Wnt are inhibited (Fig. 3E). Examining expression *hoxc10, hoxd11,* and *hoxa13* at 24hpa, we find that normally these transcripts are abundant in the regenerating axial tissue but that they fail to be induced when tadpoles are treated with 2ME, Ech, or IWR (Fig. 3F). These results suggest that *hox* genes are activated in response to injury and that this activation depends on both Hif1α and Wnt.

**Figure 3:**
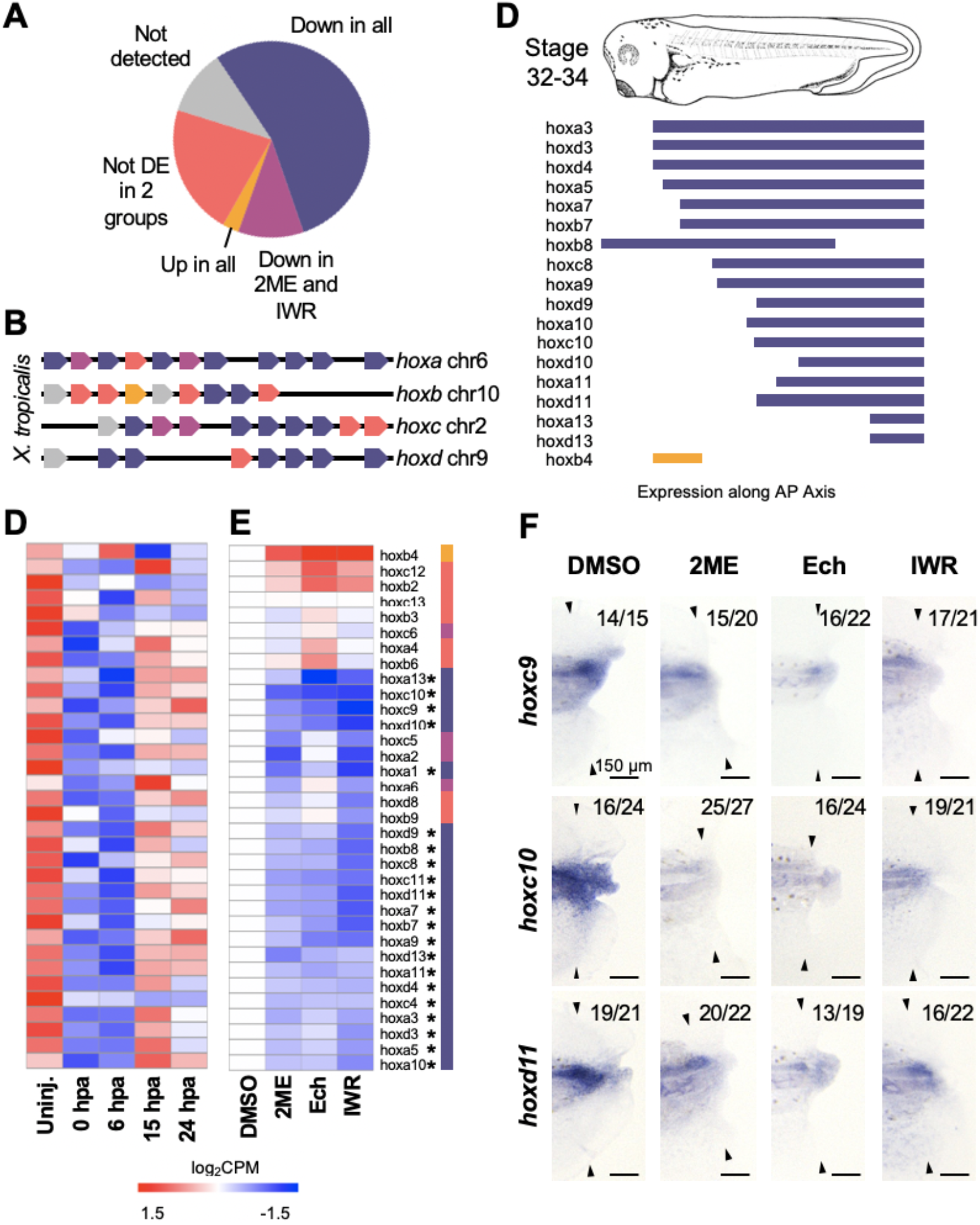
Posterior *hox* gene expression in regeneration requires Hif1α and Wnt. A) Pie chart of *hox* gene expression changes following 2ME, Ech, or IWR treatment. B) Schematic representation of *hox* gene organization in *Xenopus.* Color coding is as in (A). C) Schematic depicting *hox* gene expression in tailbud stage tadpoles. Color coding is as in (A). D) Heatmap displaying all detected *hox* genes across a regeneration timecourse. E) Heatmap displaying all detected *hox* genes clustered by expression. Genes significantly differentially expressed in all treatments are indicated by *. F) *in situ* hybridization for selected *hox* genes at 24hpa following treatment with DMSO, 2ME, Ech, or IWR. Arrowheads indicate amputation plane. Scale bars in E are 150 μm.

### Wnt signaling ligands and receptors are not regulated by Hif1α

Because the expression of posterior patterning genes was decreased if either Hif1α or Wnt signaling was blocked, we asked if Hif1α might be acting upstream of Wnt signaling, such that inhibition of Hif1α caused an indirect downregulation of Wnt signaling. Specifically, we asked whether inhibition of Hif1α resulted in downregulation of Wnt ligands or receptors. To test this, we examined expression of Wnt ligands and receptors and found that the majority were not differentially expressed following Hif1α or Wnt inhibition, several had an increase in expression under these treatments, and only *wnt5a* and *wnt5b* are weakly downregulated (Fig. 4A). This suggests to us that it is unlikely that the primary role of Hif1α in patterning is to transcriptionally upregulate Wnt ligands or receptors. We also considered that Hif1α might be required to repress expression of components of the β-catenin degradation complex but found that *dsh2* and *axin2* expression are decreased upon Hif1α inhibition, while other components are not significantly affected (Fig. 4A). These results suggest that the sensitivity of Wnt target genes to Hif1α perturbation is less likely due to Hif1α being a direct activator of Wnt signaling components or repressor of factors that destabilize β-catenin. The upregulation of several Wnt signaling components that we observe following 2ME, Ech, and IWR treatments does suggest to us that there may be compensatory upregulation of these factors when Wnt or Hif1α is inhibited, but it is not clear that this regulation is direct.

**Figure 4:**
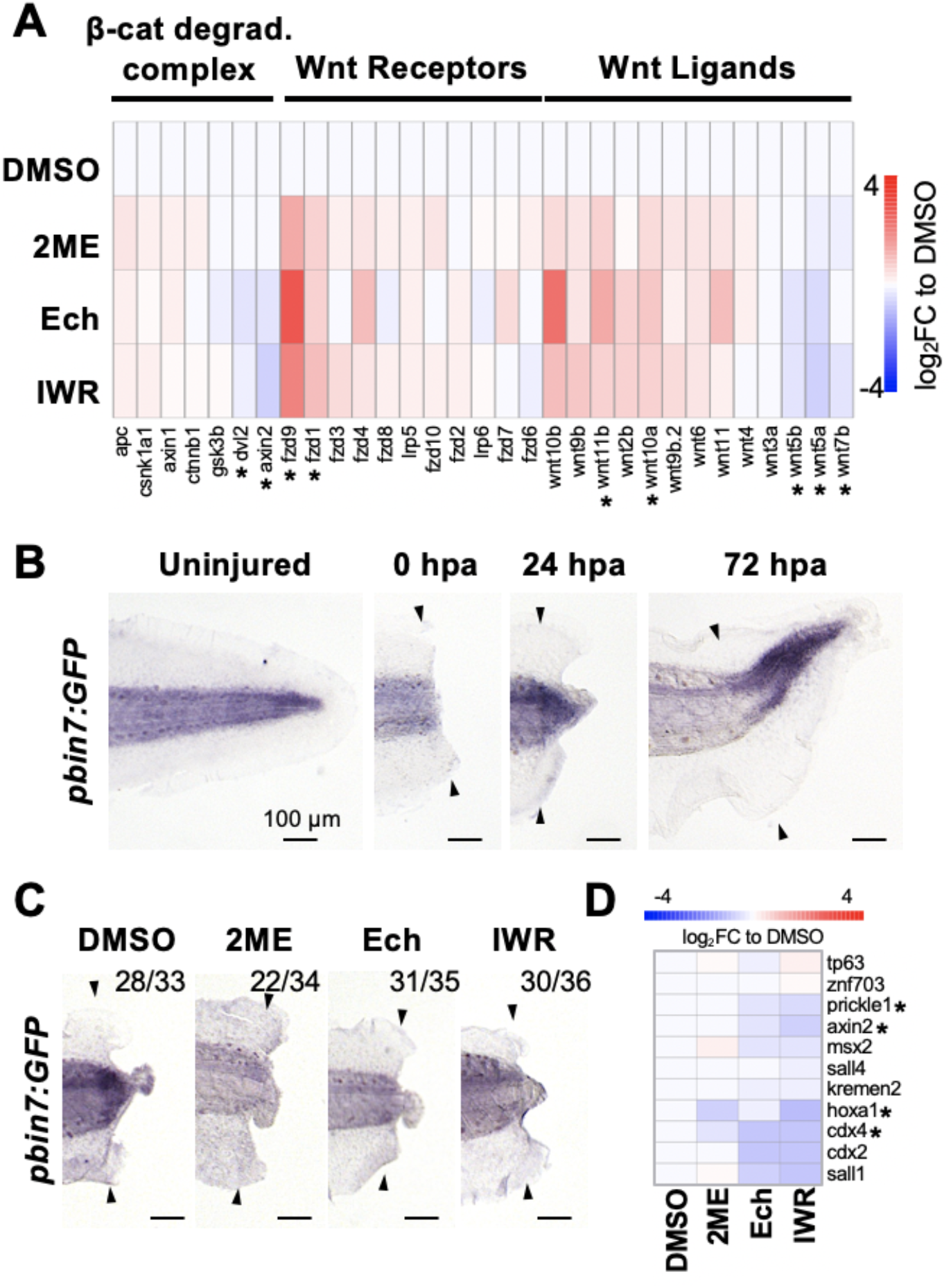
Hif1α directs WRE activity in regeneration. A) Heatmap displaying genes of the Wnt signaling pathway ordered by expression in IWR and sorted by role in pathway. Genes differentially expressed in all treatments are indicated by *. B) Timecourse of WRE-driven GFP transcript in *pbin7:GFP* tadpoles via *in situ hybridization.* C) Visualization of GFP transcripts at 24hpa following treatment with DMSO, 2ME, Ech, or IWR. D) Heatmap displaying posterior neural patterning Wnt target genes. Genes differentially expressed in all treatments are indicated by *. Indicated numbers in C represent number of tails with prevented phenotype over total number assayed. Scale bars in B-C are 150 μm. Arrowheads indicate amputation plane.

### HIf1a is required for expression of Wnt responsive elements and direct target genes of canonical Wnt signaling

Our data to this point suggested that Hif1α is required for expression of many of the same patterning genes targeted by canonical Wnt signaling, but that it is not likely to act by directly upregulating Wnt ligands or receptors. We next asked more specifically whether Hif1α acts upstream of Wnt-dependent gene expression by asking if Hif1α perturbation affects expression of Wnt-responsive promoter elements (WREs). To visualize when and where Wnt-dependent transcription is active during regeneration, we utilized the *pbin7:GFP* transgenic line, in which GFP expression serves as a readout of Wnt activity via WREs (Tran & Vleminckx, 2014). Although GFP fluorescence in the regenerating tail was relatively faint and variable, by using *in situ* hybridization to visualize GFP transcripts directly, we found *pbin7:GFP* transcripts throughout the axial tissues in uninjured tails which appears to be strongly upregulated in the posterior extreme of the tail (Fig. 4B, Supp. Fig. 3). Following amputation, we find GFP transcript localized to regenerating tissue 24hpa and find that this transcription is sustained until 72hpa (Fig. 4B). To confirm that *pbin7:GFP* transcripts are a reliable readout of Wnt activity, we treated regenerating tadpoles from this line with IWR, and found that IWR treatment reduced GFP expression at 24hpa (Fig. 4C). Inhibition of Hif1α with either 2ME or Ech also reduced GFP expression at 24hpa, at a comparable degree to IWR (Fig. 4C). As an independent test of whether Hif1α is required for Wnt-dependent gene expression, we examined expression of established direct target genes of canonical Wnt signaling established at gastrula and neurula stages (Kjolby & Harland, 2017; Young et al., 2014). We find that, while not all of these targets are still sensitive to Wnt inhibition in regeneration, those that are downregulated by Wnt inhibition, including well-established targets such as *axin2, cdx2, cdx4, prickle, sall1* and *sall4,* are also downregulated by Hif1a inhibition (Fig. 4D). These results suggest that activation of regeneration specific Wnt target requires Hif1α as well as Wnt.

### Hif1α regulates anteroposterior patterning during neurula stages

Having seen that Hif1α is required during regeneration to activate expression of posterior neural Wnt targets and the WRE during regeneration, we asked if Hif1α modulated Wnt-mediated gene expression in other contexts. To do this, we blocked Hif1α translation using a translation blocking morpholino (MO) and assayed *pbin7:GFP* expression during embryonic neural patterning. GFP expression at stage 18 spans the posterior of the neural folds with a clear anterior boundary in agreement with previous descriptions and with the well-known function of Wnt signaling in posteriorization of the neural tube (Supp. Fig. 4) (Borday et al., 2018; Kiecker & Niehrs, 2001). As previously reported, Hif1α inhibition via MO reduced neural crest migration as visualized with *snai2* (Barriga et al., 2013). Embryos in which Hif1α MO was injected unilaterally into 1 cell of the embryo at the 2-cell stage, showed posteriorly-shifted *pbin7:GFP* expression on the injected side (Supp. Fig. 4). This reflects an anteriorization of that side, as would be expected for loss of Wnt-dependent transcriptional activation. To confirm this, we assayed expression of regional neural tube markers (*sox2*, *en2*, *egr2*, *cdx4*) in unilaterally injected Hif1α morphants at stage 20 (Bradley et al., 1993; Brivanlou & Harland, 1989; Mancilla & Mayor, 1996; Northrop & Kimelman, 1994). *Sox2,* a pan-neural marker, is largely unaffected by Hif1α MO, suggesting that Hif1α is not required for neural induction. A marker of the forebrain, *otx2,* reveals an expansion of this anterior domain following Hif1α antagonism. The hindbrain marker, *en2,* and rhombomere 3 and 5 marker, *egr2*, are shifted towards the posterior following Hif1α MO injection. The posterior spinal cord marker, *cdx4* shows a slight reduction in its anterior boundary (Supp. Fig. 4). These results suggest that Hif1α is necessary for establishing posterior identity of the embryo during neurulation as well as a regenerative context.

## Discussion

Regenerative healing requires that initial wounding signals of injury be coupled not to scarring, but to growth and patterning of new tissues. Injury-induced stresses, including ROS, bioelectrical signals, and innate immune cell recruitment, are critical to regeneration across taxa (Kakebeen & Wills, 2019; Tseng & Levin, 2008). Similarly, developmental patterning signals including Wnt, FGF, BMP, and TGF-β are required for full tail regeneration in *Xenopus* (Beck et al., 2003; Christen et al., 2003; Lin & Slack, 2008). In some cases, injury stresses are known to act upstream of specific growth factor pathways, as in the case of ROS, which acts upstream of FGF in *X. laevis* tail regeneration (Love et al., 2013). Still, the mechanistic link between these aspects of regeneration has been unclear. Recently, Hif1α has been shown to be required for regeneration in *X. laevis*, acting in the first hour after injury and responding to injury-induced hypoxia and, at least indirectly, ROS (Ferreira et al., 2018). Here we propose that Hif1α bridges injury to transcriptional remodeling during regeneration by activating a broad suite of posterior patterning factors and regulatory elements that are also targeted by Wnt signaling.

Interactions between Hif1α and Wnt signaling have been described in several *in vitro* or *ex vivo* models. In prostate cancer cell lines and embryoid bodies, Hif1α has been shown to increase β-catenin target gene expression to drive differentiation (Luo et al., 2018; Večeřa et al., 2017). Hif1α has also been reported to be essential for transcription via Wnt-responsive elements in colon cancer cell lines (Rohwer et al., 2019). An *in vivo* model of murine muscle regeneration found that Hif1α actually represses Wnt signaling to inhibit regeneration (Majmundar et al., 2015). Our study demonstrates that Hif1α is required to activate Wnt target genes in the context of the whole organism, during both axial regeneration and early embryonic patterning.

One of the principal findings of our analysis is that the majority of *hox* genes are coordinately regulated by Hif1α and Wnt in our analysis. Of note, *hox* gene expression in *Xenopus* does not follow typical temporal co-linearity (Kondo et al., 2017) and we found that posterior *hox* expression domains are largely overlapping in the tail (Fig. 3E). Previous work has found that several *hox* genes, including *hoxc10* and *hoxd13,* are activated in regenerating limbs and tails in *Xenopus,* and Hoxc13 orthologs are also required for regeneration in the zebrafish tail fin (Thummel et al., 2007). It has also been proposed that ectopic *hox* gene expression many contribute to homeotic transformation of regenerating tails into limbs in *Rana temporaria* via retinyl palmitate (Christen et al., 2003; Maden, 1993). These observations could point to a mechanism in which regenerating tissues re-establish positional identity through *hox* gene expression, and our results suggest that Hif1α and Wnt are both required for this positional mapping in the regenerating tail. Our data further support that Hif1α and Wnt coordinately regulate other posteriorizing genes in regeneration, including FGF, *cdx*, and *sal* family members.

We find that Hif1α depletion anteriorizes the neural tube, suggesting that tail regeneration could be recapitulating similar gene expression programs as embryonic regionalization. Our study does not preclude the possibility that other developmental pathways, such as FGF and retinoic acid signaling, are also involved in establishing posterior identify, but we suggest that joint regulation by Hif1α and Wnt help drive this process.

The specific mechanism by which Hif1α and Wnt signaling converge on shared gene targets is not yet clear, and our work is consistent with several possibilities. While our RNASeq data suggest that Hif1α inhibition does not lead to transcriptional downregulation of most Wnt ligands, receptors, or effector proteins, it remains possible that Hif1α could be contributing to Wnt target regulation through alterations in the stability, localization, or longevity of any of these components. Our finding that Hif1α is required for TCF/LEF reporter expression suggests that Hif1a does have a specific effect on transcriptional regulation at Wnt reporter elements. However, this could be mediated by direct binding of Hif1α at these sites, as is the case in colon cancer (Rohwer et al., 2019), or by Hif1α forming a complex with β-catenin or TCF, or by indirectly enhancing the ability of β-catenin to activate these sites. We are eager to pursue these possibilities further.

Our study expands the range of processes and genes known to be targeted by Hif1α in Xenopus, where it was previously known to regulate heart development, retinal progenitor proliferation and redox balance after injury (Barriga et al., 2013; Ferreira et al., 2018; Nagao et al., 2008). That Hif1α is capable of interacting with Wnt signaling is not a new premise, but here we show for the first time that Hif1α can activate Wnt-dependent gene expression during vertebrate regeneration as well as embryonic neural pattering. Further, we demonstrate that Hif1α is essential for establishing cell fates in the regenerating tail. As we find that Hif1α directs posteriorization during regeneration and development via Wnt target genes, we are left with the possibility that many other Wnt dependent processes may be regulated by Hif1α, providing another level of regulation to the establishment of embryonic gradients and cell fate branches.

## Acknowledgements

We thank members of the Wills lab and Dana Miller (University of Washington) for thoughtful discussion throughout the course of this work. We thank Mustafa Khokha’s group for gifting the *pbin7:GFP* transgenics used in this work and Richard Harland’s group for 12/101 antibody and plasmids for *in situ* probes. We acknowledge Xenbase (www.xenbase.org) for staging, gene expression resources, and genomic reference material consulted throughout this work. JHP was supported by the Cellular and Molecular Biology Training Grant PHS NRSA T32GM007270 from NIGMS as well as National Science Foundation Graduate Research Fellowship under Grant No. DGE-1762114. This work was supported by R01NS199024 from NINDS to AEW.

## Competing Interests

The authors declare no competing interests.

## Data Availability

All sequencing data associated with this manuscript will be made publicly available on

GEO prior to final publication.

## METHODS

### *Xenopus tropicalis* husbandry and use

Use of *Xenopus tropicalis* was carried out under the approval and oversight of the IACUC committee at UW, an AALAC-accredited institution, under animal protocol 4374-01. Ovulation of adult *Xenopus tropicalis* and generation of embryos by natural matings were performed according to published methods (Khokha et al., 2002; Sive et al., 2000). Embryos were reared as described in (Khokha et al., 2002). Staging was assessed by the Nieuwkoop and Faber (NF) staging series (Nieuwkoop & Faber, 1994).

### *Xenopus tropicalis* amputation assay

NF stage 41 tadpoles were anesthetized with 0.05% ms-222 in 1/9x MR and tested for response to touch prior to amputation surgery. Once fully anesthetized, a sterilized scalpel was used to amputate the posterior third of the tail. Amputated tadpoles were removed from anesthetic media within 10 minutes of amputation into new 1/9x MR. Tadpoles were kept at a density of no more than 2.5 tadpoles per mL.

### Pharmacological inhibition

2-methoxyestradiol (Sigma, m6383-5mg) was resuspended to a 10 mM stock in DMSO, Echinomycin (Calbiochem 512-64-1) to a 0.5 mM stock in DMSO, and IWR (Sigma I0161) 10 mM stock in DMSO. Uninjured and injured tadpoles were reared with 0.1% DMSO, 5μM 2ME, 0.5μM Echinomycin, or 10μM IWR diluted in 1/9x MR until collection at 24- or 72-hours following treatment. Other concentrations were used in establishing doses and are reported in *Supp. Fig. S1.*

### Morpholino injections

Hif1α morpholino (MO) was ordered from GeneTools (sequence: CTCGCTACTACAGATCCCTCCATGC). Fertilized eggs from wild type and *pbin7:GFP* matings were de-jellied in 3% cysteine in H2O for 10-15 minutes. 10 ng of MO were injected into 1 cell at the 2-cell stage to generate unilateral morphants. Embryos were reared to NF stage 18 prior to fixation.

### Tadpole size and regeneration length measurement

Stereoscope imaging was performed on a Leica M205 FA with a color camera. Fixed tadpoles were imaged in PBS on a 1% agarose pads and measurements were recorded using the LAS software.

### Immunohistochemistry

*Xenopus tropicalis* tadpoles were fixed for 1 hour in 1x MEM with 3.7% formaldehyde at room temperature. Tadpoles were permeabilized by washing 3×15 minutes in PBS + 0.01% Triton x-100 (PBT). Tadpoles were blocked for 1 hour at room temperature in 10% CAS-block (Invitrogen #00-8120) in PBT. Then tadpoles were incubated in primary antibody [1:1 mouse anti-12/101, gift from Richard Harland (Kintner & Brockes, 1984); 1:50 mouse anti-neurofilament associated protein (DSHB 3A10) (Dodd & Jessell, 1988)] diluted in 100% CAS-block overnight at 4°C. Tadpoles were then washed 3×10 minutes at room temperature in PBT and blocked for 30 minutes in 10% CAS-block in PBT. Secondary antibody (goat antimouse 488, ThermoFisher A11001) were diluted 1:500 in 100% CAS-block and incubated for 2 hours at room temperature. Tadpoles were then washed 3×10 minutes in PBT followed by a 10-minute incubation in 1:2000 DAPI (Sigma D9542) before being washed with 1xPBS for 10 more minutes. Isolated tails were mounted on slides in ProLong Gold (ThermoFisher P36930). Images were acquired using a Leica DM 5500 B microscope using a 10X objective and processed using FIJI image analysis software (Schindelin et al., 2012).

### Whole mount *in situ* hybridization

Embryos and tadpoles were fixed overnight in 1x MEM with 3.7% formaldehyde at 4°C. *Xenopus tropicalis* multibasket *in situ* hybridization protocols were followed as described in (Khokha et al., 2002), with the notable change that pre-hybridization was always performed overnight. Isolated tails were mounted on slides in ProLong Gold (ThermoFisher P36930). Mounted tails and whole tadpoles were imaged on a Leica M205 FA with a color camera. Plasmids for *sox2, otx2, en2, egr2, cdx4,* and *snai2* probes were gifted by the Harland lab. Other probes were synthesized using the following primer pairs designed against a single exon of each mRNA: *fgf20* (forward – CTTTTGGGGATTTTGGGACT, reverse – taatacgactcactatagggGGCAGTATCTGCAGGTGGA), *pbin7:GFP* (forward – CACATGAAGCAGCACGACTT, reverse – taatacgactcactatagggTGCTCAGGTAGTGGTTGTCG), *hoxc9* (forward – CCAGCTACTGCCAGACCTTC, reverse – taatacgactcactatagggTCCAATTCCGACTTGTCCTC), *hoxc10* (forward – TCAATGGAGAAGACCCCAAG, reverse – taatacgactcactatagggTTGCTTCAGCGTCAGAATTG), *hoxd11* (forward – TGCTGTCCAAGCTCTCTTGA, reverse – taatacgactcactatagggCTCTGTGCATCACCTCCTCA).

### RNA extraction and sequencing

Tadpoles were anesthetized with 0.05% ms-222 in 1/9x MR and tested for response to touch prior to tissue collection. For 0hpa collections, a sterilized scalpel was used to amputate the posterior third of the tail followed by amputation 250 μm anterior to the wound. For 24hpa tails, 250 μm of tissue, including the regenerating tissue, was collected. 25 tails were collected for each condition and experiments were carried out in triplicate. DNA/RNA were purified by lysing tissues followed by phenol:chloroform extraction with ethanol precipitation. DNAseI treatment (Invitrogen, 18068015) was used to remove DNA. Total RNA samples were quantified using Qubit 2.0 Fluorometer (Life Technologies) and RNA integrity was checked using Agilent TapeStation 4200 (Agilent Technologies). RNA sequencing libraries were prepared using the NEBNext Ultra RNA Library Prep Kit for Illumina following manufacturer’s instructions (NEB). The sequencing libraries were validated on the Agilent TapeStation (Agilent Technologies), and quantified by using Qubit 2.0 Fluorometer (Invitrogen) as well as by quantitative PCR (KAPA Biosystems). The sequencing libraries were clustered on a single lane of a flowcell. After clustering, the flowcell was loaded on the Illumina HiSeq instrument (4000 or equivalent) according to manufacturer’s instructions. The samples were sequenced using a 2×150bp Paired End (PE) configuration. Image analysis and base calling were conducted by the HiSeq Control Software (HCS). Raw sequence data (.bcl files) generated from Illumina HiSeq was converted into fastq files and de-multiplexed using Illumina’s bcl2fastq 2.17 software. One mismatch was allowed for index sequence identification.

### Trimming, alignment, and counts

TrimGalore-0.6.5 was used to remove low quality reads (Phred33) and trim adapter sequences (http://www.bioinformatics.babraham.ac.uk/projects/trim_galore/). Pseudoalignment and count quantification was performed by Kallisto using an index built from the *Xenopus tropicalis* 9.1 transcriptome (Bray et al., 2016), returning .tsv files. These .tsv files were read into R, estimated counts for each gene were converted back to raw counts to generate a counts table suitable for processing with EdgeR.

### Multidimensional Scaling and differential expression analysis

Analysis was performed using EdgeR (Robinson et al., 2010). The counts table was made into a DGEList object and filtered for transcripts with low counts and scaled with the calcNormFactors command. MDS plots were generated using plotMDS. Differential expression between 0hpa and DMSO (24hpa), and DMSO and each treatment group – 2ME, Ech, and IWR – were performed using glmQLFTest. To determine significance, p-values for each gene generated for the DMSO vs 2ME, Ech, and IWR conditions were ordered and corrected using the Benjamini-Hochberg procedure to determine a false discovery rate (FDR). Genes considered differentially expressed between DMSO and each condition has FDR < 0.05 (as shown in *Supp. Figure 2).* Significantly downregulated genes were called using filters of FDR < 0.05 and log_2_FC < −0.2.

### Heatmap generation

Heatmaps were generated by pheatmap (Kolde, 2019). Sequencing counts were converted to counts per million (cpm) and averaged across triplicates. The average cpm was normalized to DMSO to visualize fold change between DMSO and each treatment.

### PANTHER Gene Ontology

Gene ontology was performed on the list of genes IDs called as significantly downregulated. This list was supplied to the PANTHER (Mi et al., 2019; Thomas, 2003) online portal using the reference genome for *Xenopus tropicalis* and a statistical overrepresentation test for GO biological processes was performed.

### Plotting and statistical analysis

Boxplots and stacked bar plots were generated using the R package ggplot2 (Wickham, 2009). Venn diagrams were generated using eulerr (Larsson, 2020). Length measurements were compared using ANOVA and post hoc Tukey HSD to identify differences between groups. Difference in distribution of phenotypes across multiple treatments was assessed using a chi-square test. Statistical analysis was performed in R (R Core Team, 2020).

### Analysis of previously published datasets

The heatmap in *Figure 3C* was generated as above using counts per million calculated using the supplementary counts table from GSE88975 (Chang et al., 2017).

---

**Supplemental Figure S1:**
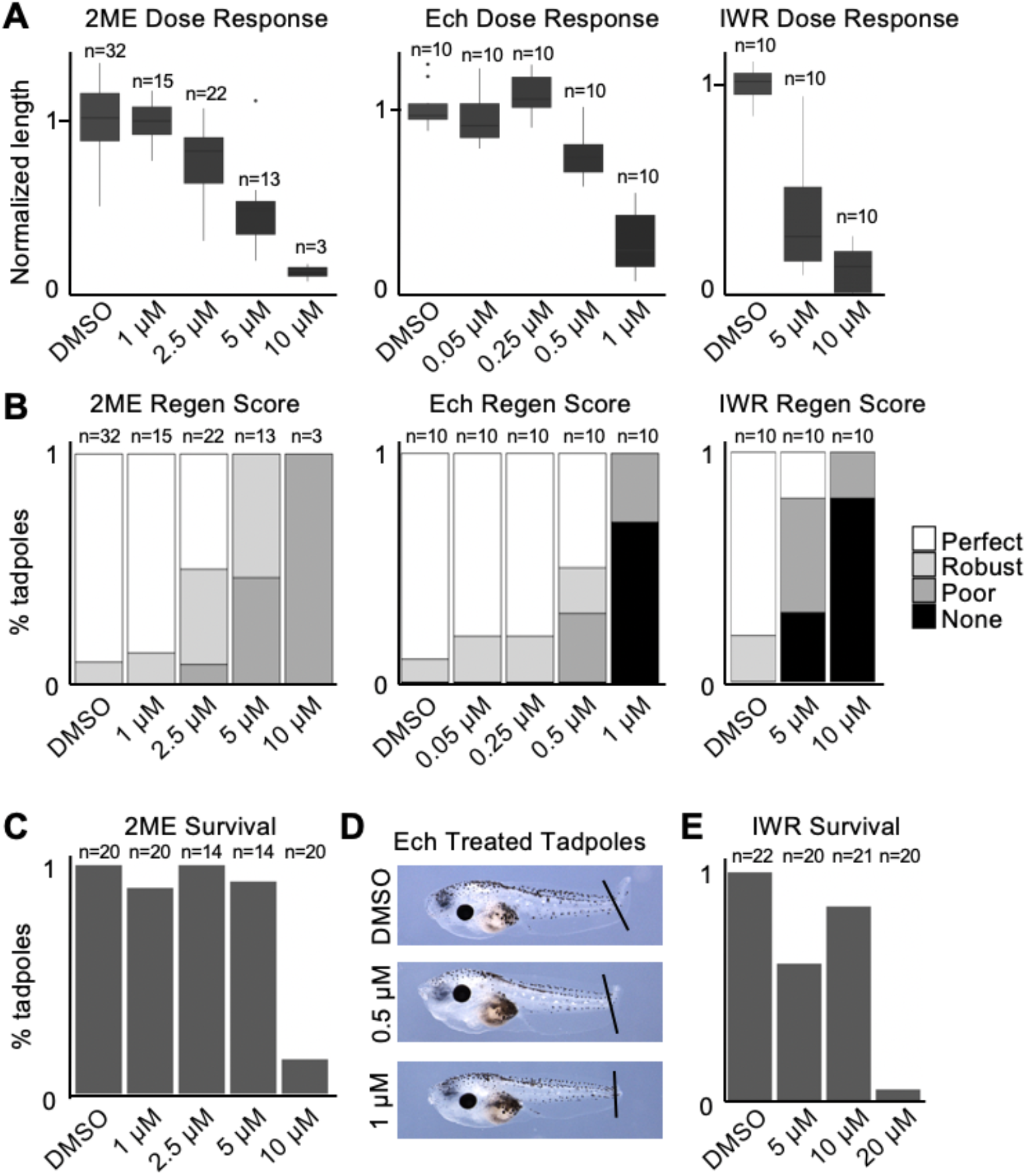
Optimization of small molecule treatments for *Xenopus tropicalis* tail regeneration. A) Quantification of regeneration length normalized to DMSO clutchmates across dose curves for 2ME, Ech, and IWR. B) Regeneration scores binned to great (full tail regeneration), good (incomplete fin regeneration), poor (very little regeneration), or none across the same tadpoles as quantified in A. C,E) Percentage of tadpoles alive at 72hpa following treatment with different concentrations of 2ME (C) and IWR (E). D) Images of tadpoles at 72hpa following treatment with DMSO or Ech. The highest concentration of Ech tested results in curling of the fin epidermis and unhealthy tadpoles.

**Supplemental Figure S2:**
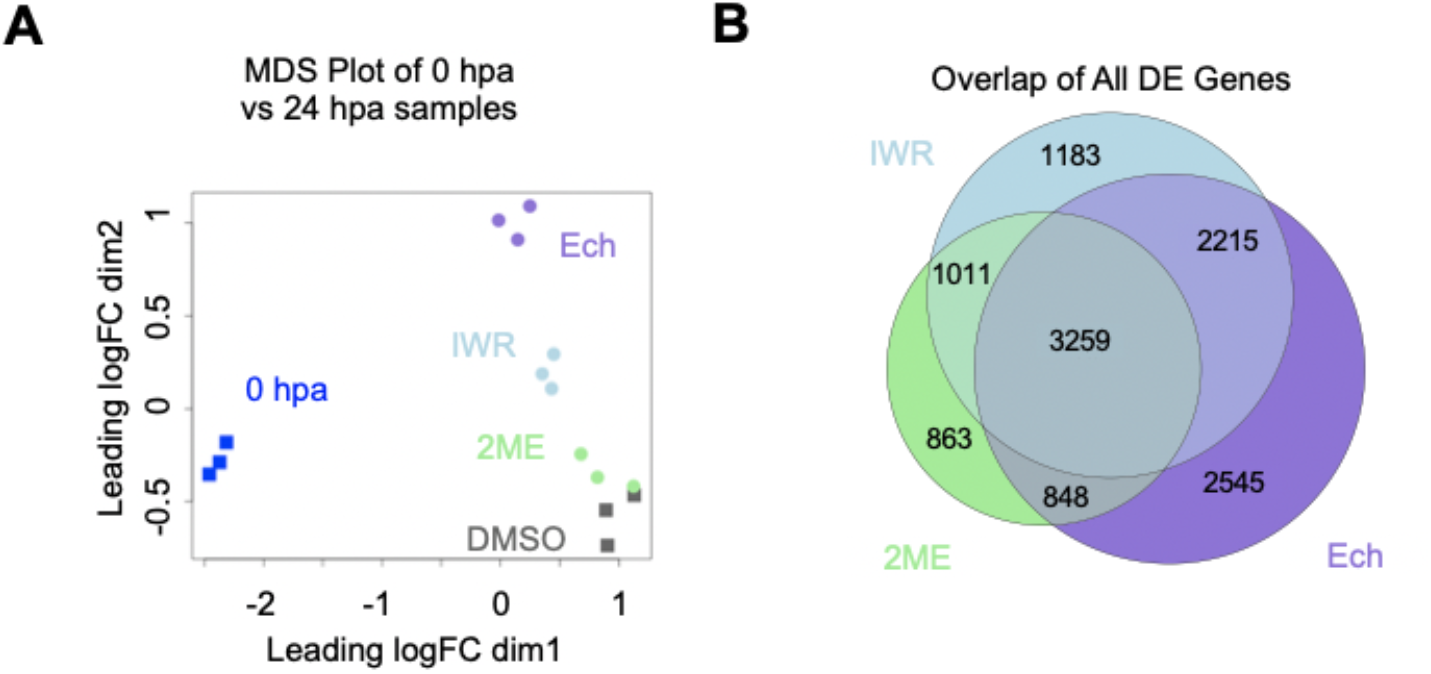
RNA-seq reveals regeneration induced changes in gene expression and Hif1α and Wnt dependent genes. A) MDS plot depicting relationship between samples including 0hpa. B) Venn diagram showing overlap in differentially expressed genes (FDR < 0.05).

**Supplemental Figure S3:**
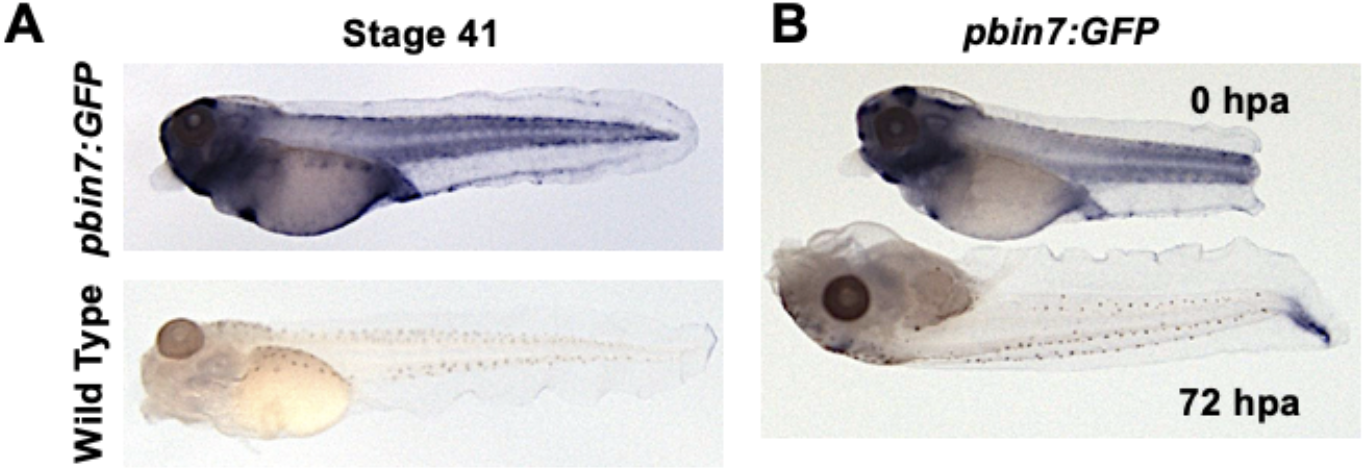
pbin7:GFP transcripts are uniquely detected in transgenic animals and axial expression declines over regenerative time. A) *In situ* hybridization for *pbin7:GFP* in WT and transgenic tadpoles. There is no detected transcript in the WT while the transgenic have patterns of expression as previously described. B) Expression in the whole tadpole from 0 to 72hpa. Axial staining decline broadly while staining in the regenerating tissue is prominently seen at 72hpa. All transgenic tadpoles can be identified at NF stage 41 by staining in the brain.

**Supplemental Figure S4:**
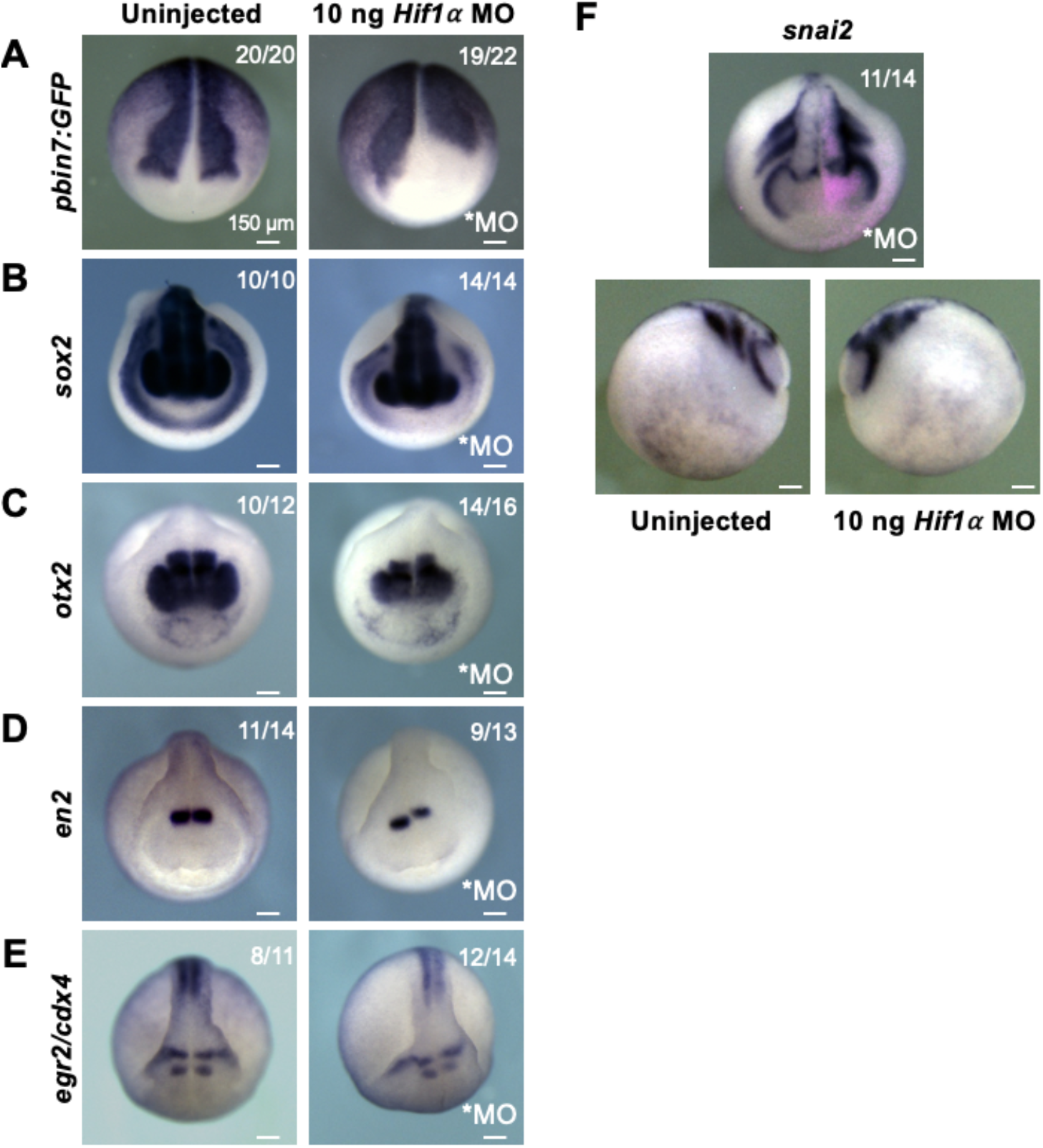
Hif1α regulates WRE expression and AP Patterning in neurulas. A-E) Dorsal views of *in situ* hybridization for *pbin7:GFP* (A), *sox2* (B), *otx2* (C), *en2*(D), *egr2* (E), and *cdx4* (E) in stage 18 (A) or stage 20 (B-E) embryos. Uninjected controls and stage matched, unilaterally injected Hif1α morphants are shown. Morpholino injected side is marked in each panel, scale. Bars are 150 μm. F) Dorsal and lateral views of i*n situ* hybridization for *snai2* in a stage 23 embryo injected unilaterally with Hif1α morpholino.

**Supplemental Table S1: GO Terms called by PANTHER from the intersection of Hif1α and Wnt downregulated genes *NOT INCLUDED IN INITIAL SUBMISSION\***

## Notes

### Competing Interest Statement

The authors have declared no competing interest.

